# A Novel Local Explainability Approach for Spectral Insight into Raw Eeg-Based Deep Learning Classifiers

**DOI:** 10.1101/2021.06.10.447983

**Authors:** Charles A. Ellis, Robyn L. Miller, Vince D. Calhoun

## Abstract

The frequency domain of electroencephalography (EEG) data has developed as a particularly important area of EEG analysis. EEG spectra have been analyzed with explainable machine learning and deep learning methods. However, as deep learning has developed, most studies use raw EEG data, which is not well-suited for traditional explainability methods. Several studies have introduced methods for spectral insight into classifiers trained on raw EEG data. These studies have provided global insight into the frequency bands that are generally important to a classifier but do not provide local insight into the frequency bands important for the classification of individual samples. This local explainability could be particularly helpful for EEG analysis domains like sleep stage classification that feature multiple evolving states. We present a novel local spectral explainability approach and use it to explain a convolutional neural network trained for automated sleep stage classification. We use our approach to show how the relative importance of different frequency bands varies over time and even within the same sleep stages. Furthermore, to better understand how our approach compares to existing methods, we compare a global estimate of spectral importance generated from our local results with an existing global spectral importance approach. We find that the δ band is most important for most sleep stages, though β is most important for the non-rapid eye movement 2 (NREM2) sleep stage. Additionally, θ is particularly important for identifying Awake and NREM1 samples. Our study represents the first approach developed for local spectral insight into deep learning classifiers trained on raw EEG time series.

## 1. INTRODUCTION

Electroencephalography (EEG) is particularly noteworthy for its rich spectral information. As such, many studies have sought to characterize how EEG frequency bands are associated with neurological disorders and function. In past decades, these studies typically used standard statistical methods to analyze spectral data, and while these statistical methods are still popular within the domain, the past 15 years have seen a growth in the use of artificial intelligence (AI) for EEG spectral analysis [1]–[5]. Many studies involving AI-based EEG spectral analysis have used explainability methods to better understand into the relative importance of different frequency bands to their topics of interest.

The majority of these studies have manually extracted spectral features. For example, some machine learning studies have used mean spectral power within frequency bands [1]–[3], and some deep learning studies have used EEG spectrograms [4], [5]. Manually extracted features offer some advantages. However, manually extracted features, by definition, limit the richness of the feature space and can limit model performance. As such, many studies have used deep learning classifiers trained on raw EEG data with the understanding that such classifiers could identify key spectral features and other relevant features in an automated manner.

Explainability for deep learning classifiers trained on raw EEG data presents problems not previously encountered in analyses involving manually extracted spectral features. Typical explainability methods can provide insight into the relative importance of each time point in raw EEG data [6]. However, knowing the importance of each time point does not necessarily imply a knowledge of the features that have been extracted and the relative importance of those features. As such, most studies using deep learning classifiers trained on raw EEG data have not integrated explainability methods.

There is a slowly growing literature surrounding the problem of spectral insight into raw EEG-based deep learning classifiers. These studies generally fall into two categories: 1) interpretable deep learning architectures and 2) post-hoc explainability methods. The interpretable deep learning architectures use filters specially designed to extract spectral features [7], [8]. However, by only extracting spectral features, these methods lose a key advantage of deep learning classifiers in that they can no longer extract non-spectral features. Several studies have developed post-hoc explainability methods to learn about the importance of different frequencies to raw EEG-based deep learning classifiers [9]–[11]. These methods are typically derivations of standard perturbation and ablation explainability methods [12] and involve modifying regions of the frequency domain and examining the resulting change in classifier performance [9]–[11]. A more recent approach used a derivation of activation maximization [11]. While these methods have great potential to positively impact deep learning-based raw EEG studies, the domain still has a key limitation.

Existing studies have focused on global, not local, explainability methods. Global methods find the importance of features to the overall model [12]. Local methods find the importance of features to the classification of individual samples [12]. Local methods could be particularly helpful for domains like automated sleep stage scoring [11] or seizure prediction [13] in which classifications are made for multiple samples across temporally evolving neurological states.

Here, we present a novel local spectral explainability approach. We provide spectral insight into a classifier trained on raw EEG data for automated sleep stage classification. We show how our approach can find the importance of different frequency bands over time. Further, to evaluate our approach within the context of existing methods, we use our local approach to form a global estimate of spectral importance and compare our results to an existing global method [10].

## 2. METHODS

Here, we discuss our dataset, preprocessing, CNN, and explainability methods.

### 2.1. Description of Data

We analyzed Sleep Cassette Data from the Sleep EDF Database [14] on Physionet [15]. The dataset consisted of 153 approximately 20-hour recordings from 78 subjects between the ages of 25 and 101 years. All but three of the subjects had 2 recordings. While the dataset included several modalities, we only used EEG data from the Fpz-Cz electrode. The EEG was recorded at a sampling frequency of 100 Hertz (Hz). Each time point was annotated by a technician as Unmarked, Movement, Awake, Rapid Eye Movement (REM), non-REM 1 (NREM1), NREM2, NREM3, or NREM4.

### 2.2. Description of Data Preprocessing

We segmented the data into 30-second segments and removed Unmarked and Movement samples. We discarded all Awake samples at the start of each recording. We removed a majority of the Awake samples starting from the end of the recording to make the number of Awake samples closer to the number of NREM2 samples. We consolidated NREM3 and NREM4 samples into a single NREM3 class [16]. After sample removal, we z-scored each recording individually. There were 215,668 total samples with 39.43%, 9.98%, 32.05%, 06.05%, and 22.98% belonging to the A, NREM1, NREM2, NREM3, and REM classes, respectively.

### 2.3. Description of Classifier

We used a preexisting CNN architecture [17] that has been used in multiple sleep stage classification studies [18], [19]. It is displayed in Figure 1. We used 10-fold cross validation to train the model with 63, 7, and 8 subjects randomly assigned to training, validation, and test sets, respectively. We used categorical cross entropy loss and weighted each class to account for class imbalances. We used an Adam optimizer with an initial learning rate of 0.001 that decreased after each 5 epochs with no improvement in validation accuracy. For each fold, we used precision, recall, and F1 score to quantify performance for each class. We calculated the mean (μ) and standard deviation (σ) of the performance across folds to summarize the overall model performance.

**Figure 1.**
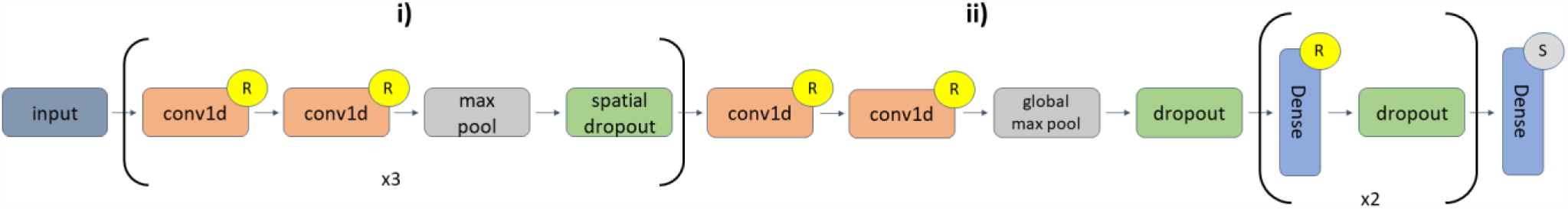
CNN Architecture. There are 6 1D convolutional (conv1d) layers in **i)**. The first repeat has two conv1d layers (number of filters = 16, kernel size = 5), a max pooling layer (pool size = 2), and a spatial dropout layer (rate = 0.01). The second repeat has two conv1d layers (number of filters = 32, kernel size = 3), a max pooling layer (pool size = 2), and a spatial dropout layer (rate = 0.01). The third repeat has 2 conv1d layers (number of filters = 32, kernel size = 3), max pooling (pool size = 2), and spatial dropout (rate = 0.01). In **ii**), the last two conv1d layers (number of filters = 256, filter size = 3) are followed by global max pooling and dropout (rate = 0.01). The first dense layer (number of nodes = 64) has a dropout layer (rate = 0.1), and the second dense layer (number of nodes = 64) has a dropout layer (rate = 0.05). The last dense layer has 5 nodes. Layers with an “R” or “S” are followed by ReLU or Softmax activation functions, respectively.

### 2.4. Description of EasyPEASI

We implemented EasyPEASI [10] as a comparison for our approach. It consists of multiple steps. (1) Calculate the classifier performance across a set of samples. (2) Convert the samples to the frequency domain. (3) Calculate the μ and σ of each spectral value. (4) Replace the values within a frequency band across all samples with values randomly generated with the previously calculated μ and σ. (5) Convert the modified samples back to the time domain. (6) Calculate the difference in classifier performance between the modified and original samples. (7) Repeat Steps 2 through 6 for each frequency band. (8) Repeat Steps 2 through 7 K times.

Our implementation differed slightly from the original implementation. We used the F1 for each class and the weighted F1 across classes as performance metrics, rather than accuracy. Also, we did not use significance testing to find key frequency bands. When converting to and from the frequency domain, we used a Fast Fourier transform (FFT) and inverse FFT. We used 5 frequency bands, not the 6 bands of the original paper. We used δ (0 – 4 Hz), θ (4 – 8 Hz), α (8–12 Hz), β (12 – 25 Hz), and γ (25 – 50 Hz). We applied EasyPEASI to the test set of each fold, rather than the training set. We perturbed the samples K = 100 times.

### 2.5. Description of Novel Local Approach

Similar to EasyPEASI, our novel approach involved converting samples into the frequency domain, perturbing the samples, and converting back to the time domain. (1) We calculated the classification probability for each sample. (2) We converted the samples into the frequency domain. (3) We substituted values with a selected frequency band from other samples in the dataset into values of a particular sample M times. (4) We calculated the percent change in classification probability from the original probability for a sample to the probabilities of the perturbed samples. (5) We repeated Steps 3-4 for each frequency band. (6) We repeated Steps 2 – 5 for each sample M times. When converting to and from the frequency domain, we used an FFT and inverse FFT. Also, we used the δ, θ, α, β, and γ bands. After computing local perturbation percent change values for each test sample, we calculated the mean of their magnitude for correctly classified samples within each fold and also displayed the local perturbation percent change values for a 2-hour period over time. There were some differences between our approach and EasyPEASI. We perturbed each sample individually. We perturbed each sample with values from other samples within the dataset. This did not require an assumption of spectra across samples having a normal distribution. In our implementation, we perturbed each sample M = 100 times.

## 3. RESULTS AND DISCUSSION

Here, we describe and discuss our classification results and the results for EasyPEASI and our approach. We then discuss next steps and study limitations.

### 3.1. Classifier Performance

Table 1 shows the μ and σ of the model performance for each class. The model had the best performance for the Awake class. This is not unexpected given that the EEG of Awake samples is so distinct from NREM samples [16] and that the Awake class had the largest number of samples. The model performance for NREM2 was less than the Awake performance, even though they had a comparable number of samples. This is possibly because the patterns differentiating Awake from other classes were more distinct. The model had the lowest performance for NREM1, similar to [18]. The model performed well for NREM2, NREM3, and REM.

**Table 1.** Classification Performance Results

|  | Awake | NREM1 | NREM2 | NREM3 | REM |
| --- | --- | --- | --- | --- | --- |
| **F1** | 91.28±02.53 | 41.94±03.07 | 77.38±03.32 | 66.59±11.32 | 68.71±05.97 |
| **Precision** | 96.82±01.44 | 36.10±05.23 | 84.97±04.09 | 57.45±11.33 | 64.88±10.75 |
| **Recall** | 86.45±04.27 | 52.04±08.26 | 71.66±07.23 | 80.66±14.96 | 74.57±02.58 |

### 3.2. EasyPEASI and Global Importance Estimate with Local Perturbation Approach

Figure 2 shows the results for EasyPEASI across all 10 folds. Based upon the weighted F1 score, δ, followed by θ and β, was the most important band when all classes were included. For identifying each class other than REM, δ was most important. For Awake, either θ or γ were of second greatest importance, and for NREM3, either θ or β were of second highest importance. For NREM1 and NREM2, β seemed to be of second highest importance. For REM, δ and θ were of comparable importance, though θ had a slightly larger median value. Interestingly, α seemed to be of least importance across all classes and most individual classes other than Awake.

**Figure 2.**
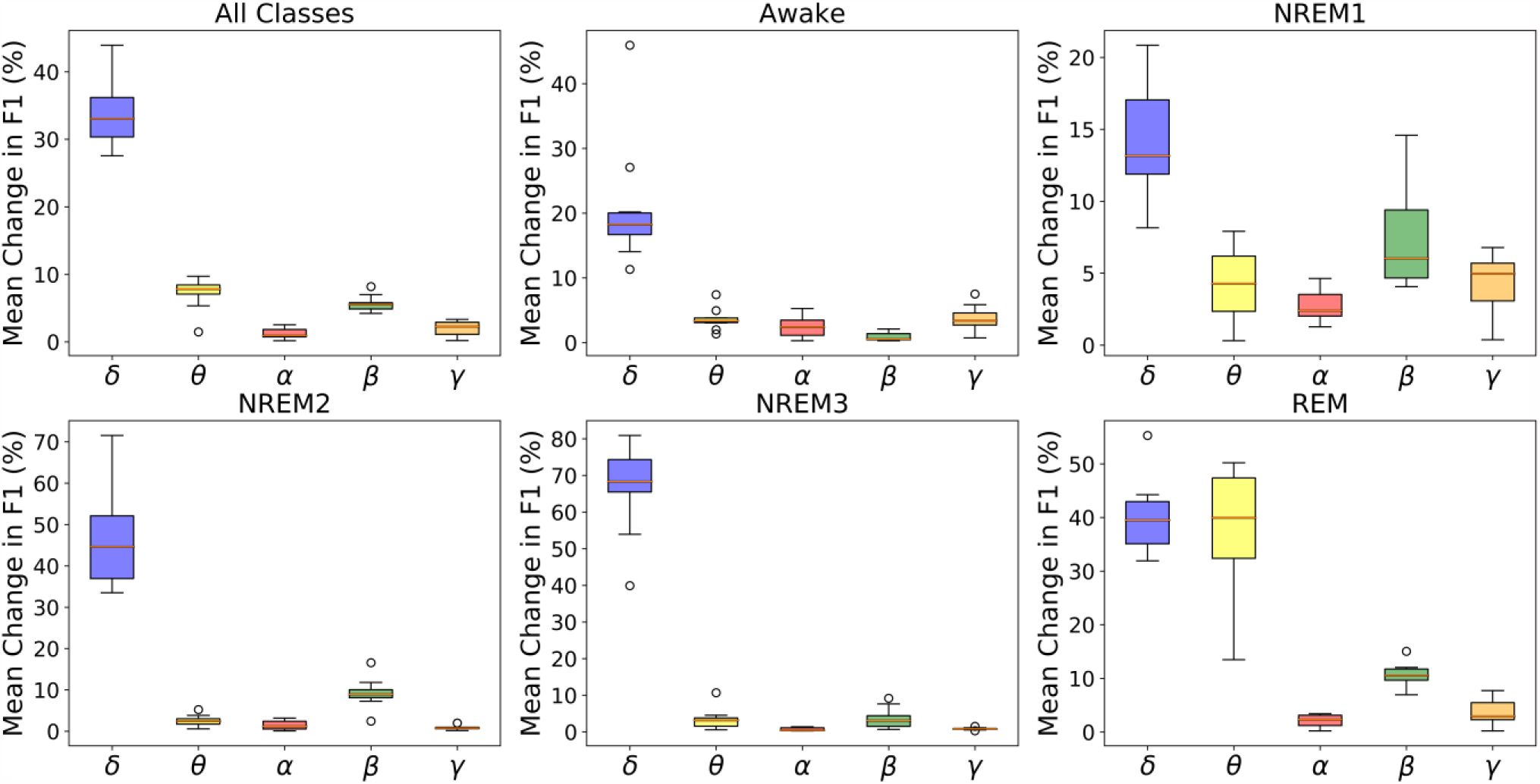
Boxplot of EasyPEASI Results. This shows the absolute change in weighted F1 across all classes following perturbation and for the change in F1 for each individual class following perturbation. The y-axis of each panel indicates the mean absolute change in F1 for the 100 perturbations within each fold. The x-axis shows the five frequency bands.

Figure 3 shows how our novel local explainability approach can estimate global importance for correctly classified samples belonging to each class. Across stages, δ was generally most important. This was the case for all sleep stages except for NREM2, in which β was most important. For Awake and NREM1, θ was of second highest importance, and for NREM3 and REM, β was of second highest importance. The importance of δ was consistent across both explainability methods. Key differences can be seen for REM, NREM1, and NREM2 classes. For EasyPEASI in REM, θ was of slightly higher mean importance than δ but was not as important for our local perturbation approach. Additionally, for NREM1, θ was of second highest importance in our approach but not nearly so important in EasyPEASI. There are several possible explanations for the differences in results. (1) EasyPEASI assumes that a particular frequency value across all samples follows a normal distribution, which may or may not be a valid assumption. (2) Our approach looks at the percent change in the classification probability for a sample. This measure can capture finer variations in the effect of a perturbed frequency band than the measure used by EasyPEASI, which requires a complete change in predicted class. Lastly, the results for both methods did not clearly conform to prior knowledge within the sleep scoring domain [16]. However, our results are very similar to those of [5] in which the authors trained a 2D-CNN for automated sleep stage classification with EEG spectrograms. This could indicate that, in some instances, deep neural networks may rely upon spectral patterns that are distinct from those used by physicians. The capacity of neural networks to learn nonlinear patterns and reliance of physicians upon linear patterns could explain this difference.

**Figure 3.**
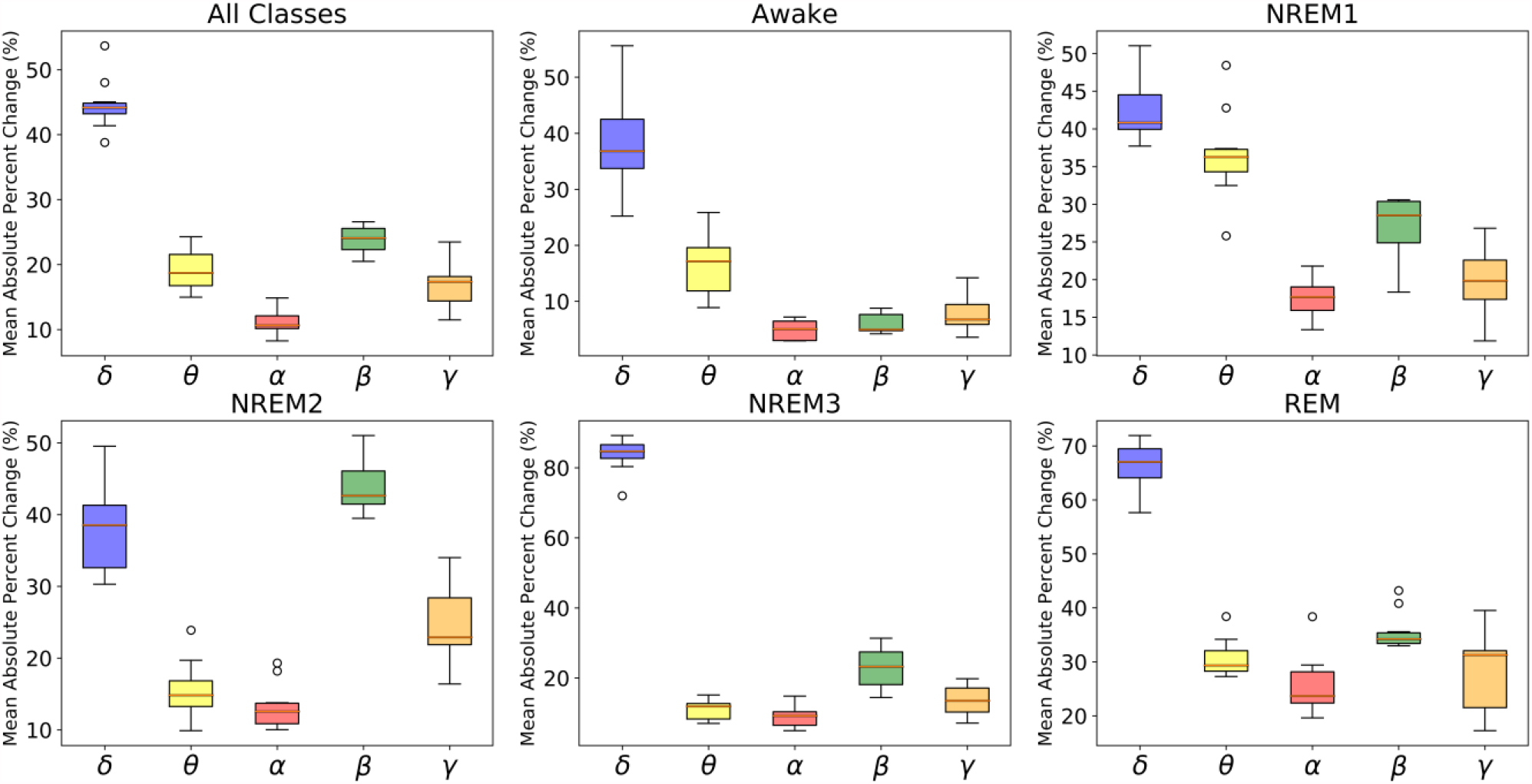
Boxplot of Local Perturbation Results. This shows the mean absolute percent change across all correctly classified samples in all classes and each individual class. The x-axis shows each of the five frequency bands.

### 3.3. Local Perturbation Over Time

Figure 4 shows the importance of each frequency band over a 2-hour recording from Subject 15. It is interesting that samples within the same class had different explanations. For example, the importance of δ for NREM2 varied from 40-80% between 20 and 30 minutes and was greater than 80% from 70 to 90 minutes. While δ appeared most prominently, β was of greater importance than δ from around 10 – 20 minutes and 40 – 60 minutes. It was also interesting how β decreased in importance from 40 to 60 minutes. It is worth noting that θ was particularly important for some NREM2 (20–40 minutes) and NREM3 (100 – 120 minutes) periods. While the results showed an importance of θ for NREM1, θ did not have importance for NREM2 and NREM3.

**Figure 4.**
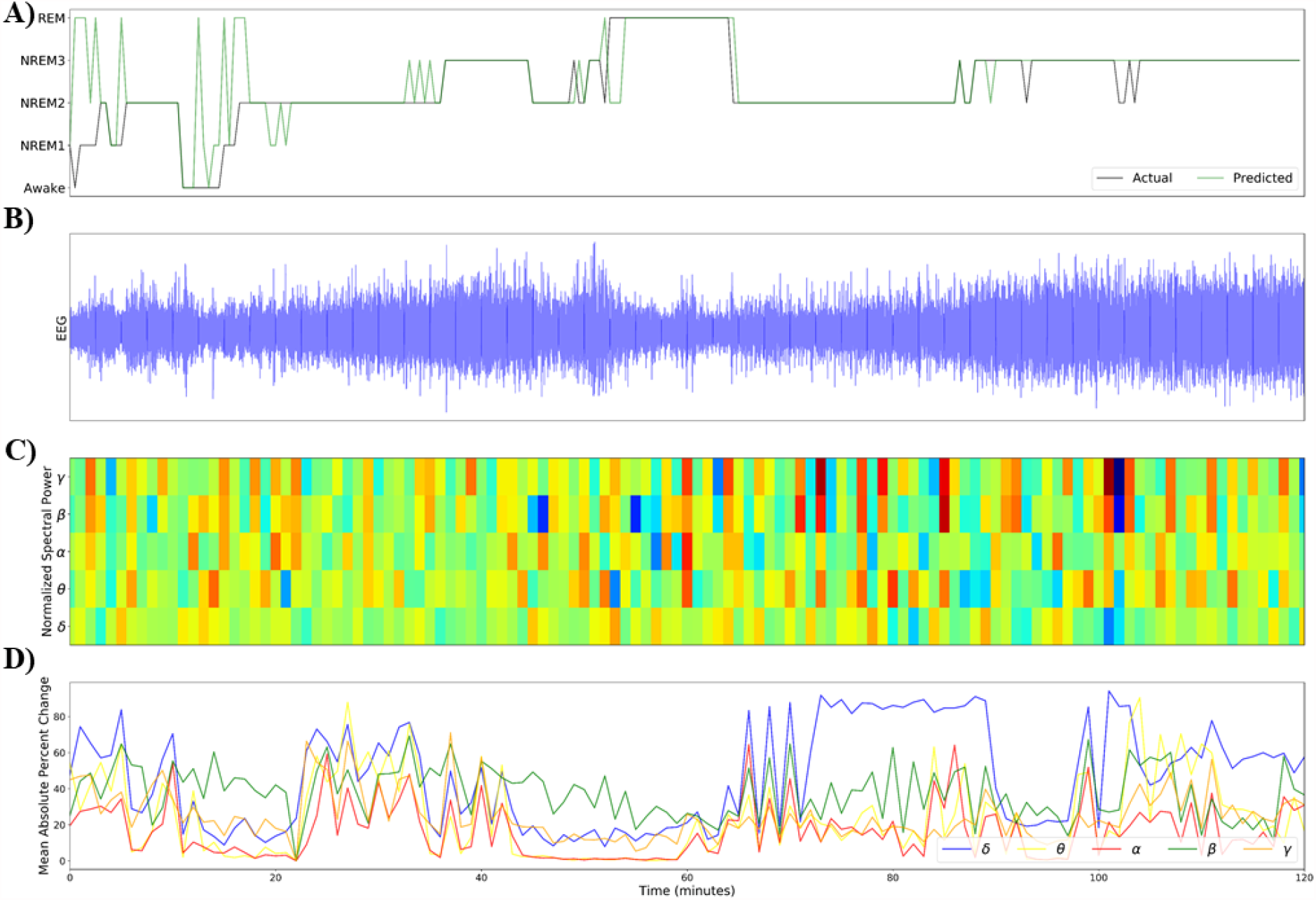
Local Spectral Explainability Results. This figure shows the local explainability results for a 2-hour window. Panel A shows the actual and predicted classes of each sample. Panel B shows EEG activity. Panel C shows the mean spectral power calculated for each 30-second sample within each frequency band. The spectral power within each band was normalized (divided by maximum spectral value within band) to enable an easier visual comparison of spectral power across bands. Panel D shows the importance for each frequency band over time. Note the legends in Panels A and D.

### 3.3. Limitations and Next Steps

It is important to note that we perturbed 30-second samples in this study. While spectral power estimates for lower frequencies are likely to be accurate for this time window, higher frequency power estimates may be inaccurate. This could have led to inaccurate spectral importance estimates for higher frequencies. While previous studies have not explored this potential problem, future studies might find it helpful to calculate spectral power for smaller segments within each sample and perturb those smaller segments simultaneously. Additionally, perturbation methods can potentially create unrealistic, out-of-distribution samples that do not accurately assess what a classifier has learned [12]. We addressed this potential problem by perturbing samples with values from within the dataset and by not assuming distribution normality like previous studies have assumed [10]. While this adaption would likely reduce the potential for out-of-distribution samples, gradient-based explainability methods avoid this problem and could potentially be applied to gain local spectral insight [20][21]. Additionally, we only presented global importance estimates for correctly classified samples. We also examined the spectral importance for incorrectly classified samples, but excluded them to enable a clearer comparison with easyPEASI. These measures could give more insight into which frequency bands may have caused misclassification. Lastly, the local explainability results could be compared with statistical analyses to determine whether demographic variables affected the model.

## 4. CONCLUSION

We presented a novel local spectral explainability approach. We showed how each frequency band affected sample classification over time. We further used our local approach to form a global estimation of the importance of different frequency bands and compared our method with easyPEASI, a global spectral explainability approach. Both methods found δ to be most important to classification. However, there were some differences in their results that could be explained by their different spectral perturbation approaches. EasyPEASI assumed that the spectral values across the dataset followed a normal distribution, which might not be a valid assumption given the number of classes and could exacerbate the out-of-distribution sample problem inherent to perturbation approaches. Our approach did not assume normality and sought to maintain the sample distribution. To the best of our knowledge, our study represents the first approach for local spectral insight into deep learning classifiers trained on raw EEG time series.

## 10. ACKNOWLEDGEMENTS

We thank Darwin A. Carbajal for helping develop Figure 1.

## REFERENCES

[1] Y. Chen et al., “Automatic Sleep Stage Classification Based on Subthalamic Local Field Potentials HHS Public Access,” IEEE Trans Neural Syst Rehabil Eng, vol. 27, no. 2, pp. 118–128, 2019, doi: 10.1109/TNSRE.

[2] M. L. Bringas Vega et al., “An Age-Adjusted EEG Source Classifier Accurately Detects School-Aged Barbadian Children That Had Protein Energy Malnutrition in the First Year of Life,” Front. Neurosci., vol. 13, Nov. 2019, doi: 10.3389/fnins.2019.01222.

[3] H. G. Yi, Z. Xie, R. Reetzke, A. G. Dimakis, and B. Chandrasekaran, “Vowel decoding from single-trial speech-evoked electrophysiological responses: A feature-based machine learning approach,” Brain Behav., vol. 7, no. 6, Jun. 2017, doi: 10.1002/brb3.665.

[4] G. Ruffini et al., “Deep Learning With EEG Spectrograms in Rapid Eye Movement Behavior Disorder,” Front. Neurol., vol. 10, Jul. 2019, doi: 10.3389/fneur.2019.00806.

[5] A. Vilamala, K. H. Madsen, and L. K. Hansen, “Deep convolutional neural networks for interpretable analysis of EEG sleep stage scoring,” IEEE Int. Work. Mach. Learn. Signal Process. MLSP, vol. 2017-Septe, no. 659860, pp. 1–6, 2017, doi: 10.1109/MLSP.2017.8168133.

[6] I. Sturm, S. Lapuschkin, W. Samek, and K. R. Müller, “Interpretable deep neural networks for single-trial EEG classification,” J. Neurosci. Methods, vol. 274, pp. 141–145, Dec. 2016, doi: 10.1016/j.jneumeth.2016.10.008.

[7] D. Borra, S. Fantozzi, and E. Magosso, “EEG Motor Execution Decoding via Interpretable Sinc-Convolutional Neural Networks,” In Mediterranean Conference on Medical and Biological Engineering and Computing, 2019, vol. 1, pp. 1515–1525, doi: 10.1007/978-3-030-31635-8.

[8] D. Borra, S. Fantozzi, and E. Magosso, “Interpretable and lightweight convolutional neural network for EEG decoding: Application to movement execution and imagination,” Neural Networks, vol. 129, pp. 55–74, 2020, doi: 10.1016/j.neunet.2020.05.032.

[9] R. T. Schirrmeister et al., “Deep learning with convolutional neural networks for EEG decoding and visualization,” Hum. Brain Mapp., vol. 38, no. 11, pp. 5391–5420, Nov. 2017, doi: 10.1002/hbm.23730.

[10] D. O. Nahmias and K. L. Kontson, “Easy Perturbation EEG Algorithm for Spectral Importance (easyPEASI): A Simple Method to Identify Important Spectral Features of EEG in Deep Learning Models,” In Proceedings of the 26th ACM SIGKDD International Conference on Knowledge Discovery & Data Mining, Aug. 2020, pp. 2398–2406, doi: 10.1145/3394486.3403289.

[11] S. Pathak, C. Lu, S. B. Nagaraj, M. van Putten, and C. Seifert, “STQS: Interpretable multi-modal Spatial-Temporal-seQuential model for automatic Sleep scoring,” Artif. Intell. Med., vol. 114, no. January, p. 102038, 2021, doi: 10.1016/j.artmed.2021.102038.

[12] C. Molnar, Interpretable Machine Learning A Guide for Making Black Box Models Explainable, 2018th-08–14th ed. Lean Pub, 2018.

[13] K. Rasheed et al., “Machine learning for predicting epileptic seizures using EEG signals: A review,” arXiv, vol. 14, pp. 139–155, 2020.

[14] B. Kemp, A. H. Zwinderman, B. Tuk, H. A. C. Kamphuisen, and J. J. L. Oberye, “Analysis of a sleep-dependent neuronal feedback loop: the slow-wave microcontinuity of the EEG,” IEEE Trans. Biomed. Eng., vol. 47, no. 9, pp. 1185–1194, 2000, doi: 10.1109/10.867928.

[15] G. AL et al., “PhysioBank, PhysioToolkit, and PhysioNet: Components of a New Research Resource for Complex Physiologic Signals,” Circulation, vol. 101, no. 23, pp. e215–e220, 2000, [Online]. Available: http://circ.ahajournals.org/content/101/23/e215.full.

[16] C. Iber, S. Ancoli-Israel, A. L. Chesson, and S. F. Quan, “The AASM Manual for Scoring of Sleep and Associated Events: Rules, Terminology, and Technical Specifications.” 2007.

[17] M. Youness, “CVxTz/EEG\_classification: v1.0,” 2020. https://github.com/CVxTz/EEG_classification (accessed Jan. 05, 2021).

[18] C. A. Ellis, R. Zhang, D. A. Carbajal, R. L. Miller, V. D. Calhoun, and M. D. Wang, “Explainable Sleep Stage Classification with Multimodal Electrophysiology Timeseries,” bioRxiv, pp. 0–3, 2021.

[19] C. A. Ellis, D. A. Carbajal, R. Zhang, R. L. Miller, V. D. Calhoun, and M. D. Wang, “An Explainable Deep Learning Approach for Multimodal Electrophysiology Classification,” bioRxiv, pp. 12–15, 2021.

[20] S. Pathak, C. Lu, S. B. Nagaraj, M. van Putten, and C. Seifert, “STQS: Interpretable multi-modal Spatial-Temporal-seQuential model for automatic Sleep scoring,” Artif. Intell. Med., vol. 114, no. January, p. 102038, 2021, doi: 10.1016/j.artmed.2021.102038.

[21] S. Bach, A. Binder, G. Montavon, F. Klauschen, K. R. Müller, and W. Samek, “On pixel-wise explanations for non-linear classifier decisions by layer-wise relevance propagation,” PLoS One, vol. 10, no. 7, Jul. 2015, doi: 10.1371/journal.pone.0130140.

